# Characterizing tissue composition through combined analysis of single-cell morphologies and transcriptional states

**DOI:** 10.1101/2020.09.05.284539

**Authors:** Feng Bao, Yue Deng, Sen Wan, Bo Wang, Qionghai Dai, Steven J. Altschuler, Lani F. Wu

## Abstract

Advances in spatial transcriptomics technologies enable optical profiling of morphological and transcriptional modalities from the same cells within tissues. Here, we present <u>mu</u>lti-modal <u>s</u>tructured <u>e</u>mbedding (MUSE), an approach to deeply characterize tissue heterogeneity through analysis of combined image and transcriptional single-cell measurements. We demonstrate that MUSE can discover cellular subpopulations missed by either modality as well as compensate for modality-specific noise. MUSE identified biologically meaningful cellular subpopulations and stereotyped spatial patterning within heterogeneous mouse cortex brain tissues, profiled by seqFISH+ or STARmap technologies. MUSE provides a framework for combining multi-modal single-cell data to reveal deeper insights into the states, functions and organization of cells in complex biological tissues.

## Introduction

Living tissues are built from ensembles of cells in different states. Microscopy provides a classical approach to identify and characterize cell types through similarities in morphology^1–3^. Developments in single-cell transcriptomics technologies provide complementary approaches to characterize cells types through similarities in transcriptional states^4–8^. Both microscopy and single-cell transcriptomics approaches have provided deep insights into cellular state, function and organization. Recent advances in spatial transcriptomics technologies, such as spatial transcriptomics (ST)^9, 10^, sequential fluorescence *in situ* hybridization (seqFISH)^11, 12^, multiplexed error-robust fluorescence *in situ* hybridization (MERFISH)^13, 14^ and spatially-resolved transcript amplicon readout mapping (STARmap)^15^, enable simultaneous morphological and transcriptional profiling from the same single cells. Here, we explore the exciting possibility that techniques in machine learning can be used to combine information from microscopy and single-cell transcriptomics to provide deeper insights into cell-type compositions that comprise tissues.

Methods that successfully combine multi-modal information hold the promise to identify biologically meaningful subpopulations that are missed by individual modalities and provide a more detailed description of tissue cell heterogeneity (**Fig. 1a**). However, effective approaches that combine multi-modal data need to overcome several challenges. Notable are requirements that: (**requirement 1**) discriminative information in each modality should be captured in the combined data structure; and (**requirement 2**) limited information from a lower-quality modality should improve—and not reduce—the resolution of the subpopulation structure that is learned from the higher-quality modality.

**Figure 1.**
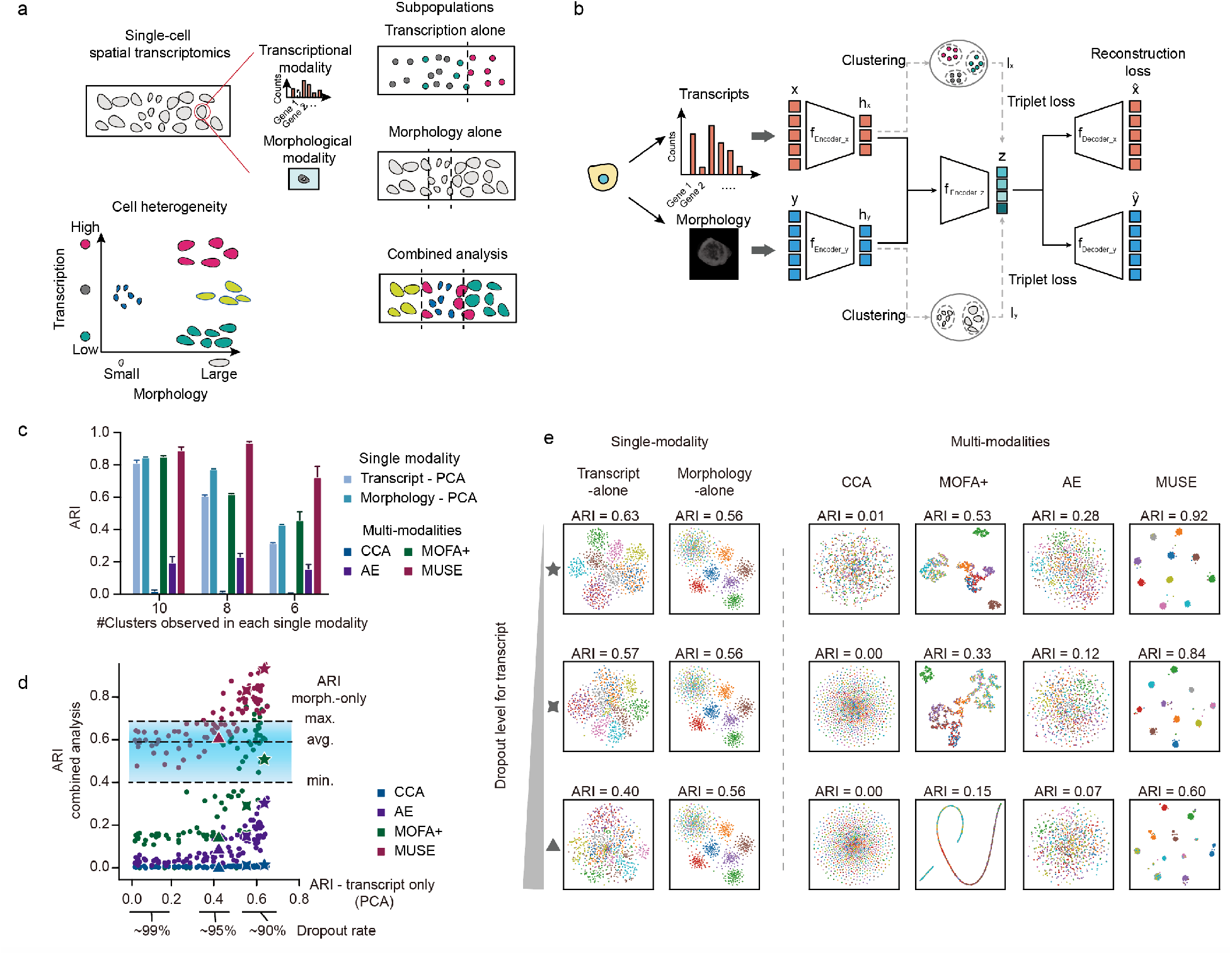
Overview of MUSE and performance evaluation on simulated data. (**a**) Cartoon indicating how single-cell morphological and transcriptional data from a tissue (rectangular slide) can be combined to reveal high-resolution characterization of tissue heterogeneity. (**b**) Overview of MUSE architecture. MUSE combines features from transcripts (x) and morphology (y) into a joint latent representation z. The reconstruction and triplet losses encourage subpopulation structure from each modality to be faithfully maintained in z. (**c-e**) Performance evaluations using simulated data. (**c**) Accuracy of identifying ground-truth high-resolution subpopulations (k=10) from lower-resolution single-modality subpopulations (k=10, 8 or 6). 1,000 cells with transcriptional and morphological profiles were simulated. Cluster accuracy is quantified using the adjusted Rand index (ARI); mean+std is shown for 10 replicates. (**d**) Accuracy of identifying ground-truth clusters over a range of dropout levels from the transcriptional modality. Dashed lines: min, ave and max ARI of morphology-modality alone. x-axis: ARI of PCA analysis on transcript-modality alone. y-axis: ARI of combined-modality methods. 3-, 4-, 5-pointed shapes: comparison of results for randomly chosen datasets, also visualized in panel (**e**). (**e**) tSNE visualizations of latent representations from single- and combined-modality methods. Colors: ground truth subpopulation labels in simulation.

Here, we present a <u>m</u>ulti-modal <u>s</u>tructured <u>e</u>mbedding (MUSE) approach that addresses these requirements. MUSE uses a deep-learning architecture to extract and integrate information from each modality into a meaningful joint representation. A self-reconstruction loss ensures that information from each modality is not lost in the process of building the joint latent representation, and a self-supervised loss ensures that phenotypic similarity of cells in each modality is preserved in the joint representation. We demonstrate these ideas first using synthetic data with known ground truth. Then we apply MUSE to two single-cell spatial transcriptomics datasets of neural cortex^12, 15^, which have the desirable properties for methods validation that many cell types have stereotyped morphologies, transcriptional biomarkers and positions within the tissue.

## Results

### MUSE architecture and training

MUSE is built on a standard multi-view autoencoder (AE) neural network architecture^16, 17^ (**Fig. 1b**). Learning is conducted in three steps: 1) **modality-specific transformations:** the input features x and y are transformed into latent representations h_x_ and h_y_; 2) **pseudo-label learning:** clustering on feature spaces h_x_ and h_y_ are performed independently to obtain pseudo-labels l_x_ and l_y_ for each modality; and 3) **joint feature learning**: the modality-specific features h_x_ and h_y_ are merged and transformed into a joint latent feature representation z. The learning process is guided by minimizing combined self-reconstruction and self-supervised loss functions. The self-reconstruction loss resembles the standard AE loss function, which encourages the learned joint feature representation (z) to faithfully retain information from the original individual input feature modalities (x and y). The self-supervised learning exploits triple-loss functions^18, 19^ to encourage cells with the same cluster label (i.e. with the same pseudo label in either l_x_ or l_y_) to remain close—and cells with different cluster labels to remain far apart—in the joint latent space. During model training, the transformation, pseudo-label learning and joint feature learning steps are iteratively performed. Model parameters in the whole neural network are jointly updated in each iteration (**Methods**). Finally, after model training, graph clustering is performed on the joint latent features (z) using PhenoGraph^20, 21^ to identify latent subpopulations (**Methods**).

### Combined analysis improves cell subpopulation identifications

To evaluate the performance of MUSE, we initially made use of simulated transcript and morphology data where ground truth subpopulation assignment for each cell and modality is known (**Methods** and **Supplementary Fig. 1**). As benchmarks, MUSE was compared with three existing approaches that enabled combining data: canonical correlation analysis (CCA)^22^, multi-omics factor analysis v2 (MOFA+)^23, 24^, and a multi-view autoencoder (AE). Results using a single modality were presented using principle component analysis (PCA) as reference. For each method, graph clustering was used to identify the underlying subpopulation structures, and the accuracy to correctly discover true cell subpopulations was quantified using the adjusted Rand index (ARI)^25^.

We first used the simulated data to assess the ability of MUSE to capture discriminative information from each modality (**requirement 1 above**). How is performance affected as the ability to discriminate subpopulations in each modality decreases? We retained 10 ground truth subpopulations in the full multi-modal space and degraded the ability of both single modalities to resolve these subpopulations by randomly merging a different group of cell cluster assignments for each modality (**Methods**). Transcriptional data were simulated using a published single-cell RNA simulator^26, 27^, and morphological features were simulated using a multi-layer neural network (**Methods**). As cluster numbers decreased, the factorization method MOFA+ maintained an accuracy level comparable to either single-modality approach while MUSE exceeded the single-modality benchmarks (**Figs. 1c**). Visualization of the latent space suggested the utility of the triplet-loss function: cells originating from the same subpopulation in either modality remained close and all true subpopulations remained well distinguished **(Supplementary Fig. 2**). How is performance affected as the number of ground-truth subpopulations increases? A potential advantage for multi-modal analysis is the ability discover more fine-grained population composition by combining heterogenous cellular properties; however, accuracies tend to decrease with increasing cellular heterogeneity. Here, we held the total number of cells constant but increased the number of subpopulations. As the number of clusters increased, CCA, AE and MOFA+ did not achieve higher ARIs than the single-modality benchmark methods; however, the guided combination of MUSE consistently outperformed single-modality methods (**Supplementary Fig. 3**).

We next assessed the performance of MUSE as data quality in one modality degrades (**requirement 2 above**). Two persistent problems in single-cell data are sequencing dropouts and noise in feature measurements^28, 29^. First, we varied dropout level for the transcript modality while leaving the simulation parameters for the morphology modality unchanged (**Methods**); as before, 10 ground-truth clusters were used. Morphology alone provided an average accuracy of ~0.6 ARI (**Fig. 1d**, horizontal dashed lines). As the dropout rate increased, the accuracy of all methods degraded, but only MUSE was able to retain the accuracy of morphology alone. Visualizing the results in latent space suggested that MUSE representations maintained a discernable subpopulation structure of 10 clusters (**Fig. 1e**). Second, we changed the noise level in both modalities, using additive Gaussian random noise with increasing variance (**Supplementary Fig. 4**). MUSE largely performed better or equal to the benchmark single-modality methods, though at extremely high noise levels the performance of MUSE and all other combined methods were strongly compromised.

Finally, we note that the choice of latent dimension had only a minimal effect on the accuracy of subpopulation identification for all compared methods (**Supplementary Fig. 5)**; further, the accuracy of MUSE was not affected by input dimension (**Supplementary Fig. 6**). While MUSE is reasonably fast for current experimental data sizes (e.g. 1000 samples in 1.5 minutes on a standard desktop, **Methods**), MUSE is slower than all other compared methods due to the inclusion of graph clustering during structured self-supervised training, (**Supplementary Fig. 7**). All simulation parameters used in experiments were summarized in **Supplementary Table 1**.

Together, these results indicated that direct combination of two feature modalities does not guarantee a better cell-subpopulation decomposition. The structured self-supervised used by MUSE enabled capturing and combining discriminative information that was not available from either modality alone. Further, MUSE was not unduly confounded by poor data quality in either one or both modalities. Thus, MUSE satisfied the two requirements we set *a priori* for a combined multi-modal method.

### MUSE analysis of mouse cortex layers (seqFISH+)

A critical challenge in assessing single-cell analysis on real data is often a lack of ground truth. However, tissues with stereotyped spatial organization of cell types can provide independent evidence to evaluate the quality of learned representations and identified subpopulations^30, 31^. A particularly good example of this is the multi-layer pattern of spatial organization in the brain cortex^32, 33^. Thus, we applied MUSE to two experimental mouse cortex datasets.

The first cortex dataset was obtained using seqFISH+ technology^12^. This dataset includes expression profiles of 10,000 genes and cell images with DAPI and Nissl staining for 523 cells. For the transcript modality, we used a standard preprocessing pipeline for scRNA and selected highly variable genes as input features x (**Methods**). For the morphological modality, we input the DAPI and Nissl images for each cell (based on the provided cell masks) into a pre-trained deep neural network (Google Inception-v3^34^) to extract morphological properties as input features y (**Methods**). We extended subpopulation analyses to include four approaches in each of three classes—using only transcriptional features x (PCA, ZIFA, SIMILR and scScope; with detailed descriptions in **Methods**), only morphological features y (PCA, MDS, Isomap and tSNE), or the combination of both x and y (CCA, MOFA+, AE and MUSE). As before, cell clusters and cluster numbers were identified automatically by performing graph clustering on the latent cell representations z (**Methods**).

Many clusters identified from each method were spatially co-localized (**Methods**; **Supplementary Figs. 8** and **9**) and showed layer-like structure (**Figs. 2a** and **b**, **Supplementary Fig. 10**). Layer-specific markers were used to assign clusters to the cortex layers, and MUSE was able to identify all four layers (L2/3, L4, L5 and L6; **Figs. 2c, d** and **Supplementary Figs. 11, 12**). By examining MUSE clusters within the same cortex layers, we observed that the morphology modality refined distinctions provided by the transcript modality (e.g. in L2/3, distinct distributions of morphological features between the subpopulations were evident; **Fig. 2e** and **Supplementary Fig. 13**). Combing morphological and transcriptional profiles provided a refined dissection of cell diversity within the cortex.

**Figure 2.**
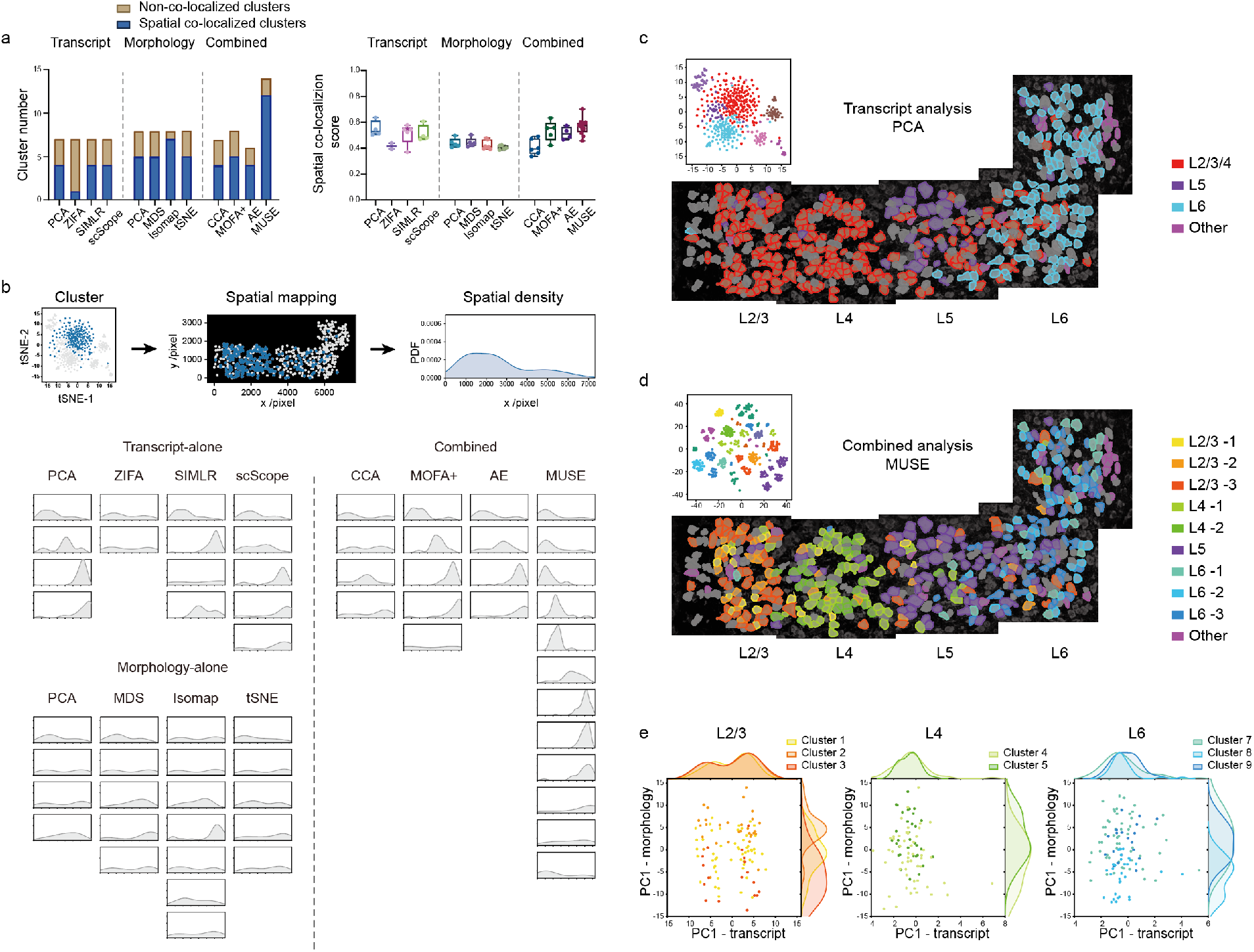
Evaluation of MUSE on seqFISH+ mouse cortex data. Analysis of cells (n=523) based on transcript (top 500 variable genes) and/or morphology (DAPI and Nissl images) modalities. (**a**) Numbers (left) and scores (right) of clusters whose cells show spatial co-localization in the tissue (**Methods**). Right box plot: median (center line), interquartile range (box) and data range (whiskers). (**b**) Visualization of spatial density in the tissue section for clusters with co-localization patterns. Top: clusters were mapped to the tissue and spatial density was quantified by kernel density estimations. Bottom: spatial density plot for each cluster. Coordinates in each subfigure are the same as the density plot in the top. (**c-d**) Spatial mapping and tSNE plots of cell clusters with co-localization patterns based on (c) transcript-only analysis using PCA or (d) MUSE. Other transcript-only methods reveal similar spatial patterns (**Supplementary Fig. 10)**. Layers were annotated using marker genes identified from differential expression analysis (**Methods**). Cell colors: consistent with PCA or MUSE in (a) (respectively). (**e**) Visualization of clusters in the same layers identified by MUSE. Plots: first principle component (PC1) of raw features from each modality. Density graphs (top, right): Gaussian kernel density estimations.

### MUSE analysis of mouse cortex layers (STARmap)

The second cortex dataset was obtained using STARmap technology. For the transcript modality, this dataset contained expression profiles of 1,020 genes; however, for the morphological modality, only cell shape masks were provided. The data were processed for the different comparison methods to obtain latent representations and subpopulations as in the previous cortex dataset (**Supplementary Fig. 14**).

We visualized the ability of different methods to “discover” cortical layer structure based on pseudo-colored cortex depth (**Fig. 3a**). SIMLR and MUSE identified the highest number of spatially co-localized clusters (**Fig. 3b**, **Supplementary Figs. 15** and **16**), and the clusters from MUSE were well separated in latent space (**Figs. 3a** and **c**). Based on anatomic annotations from the original paper, which labeled all seven layers in the cortex sample, MUSE successfully identified all neuron and non-neuron layers (**Fig. 3d**; **Methods**).

**Figure 3.**
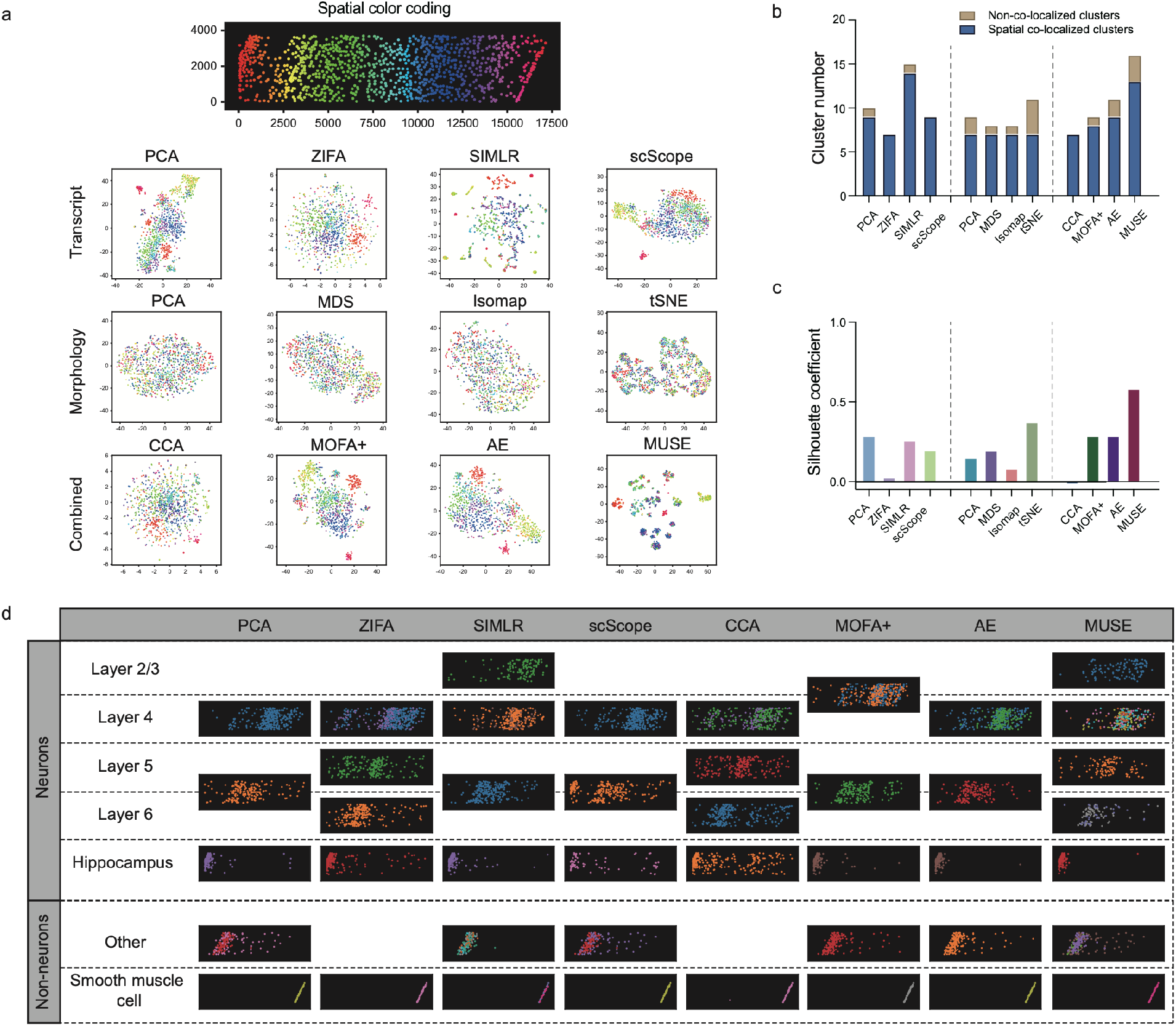
Evaluation of MUSE on STARmap cortex data. Analysis of cells (n=972) based on transcript (top 500 variable genes) and/or morphology (segmentation mask) modalities. (**a**) tSNE visualization of latent representations by different methods with pseudo-colors labeling cortex depth along x-coordinate. (**b**) Numbers of identified clusters with or without significant spatial co-localization properties. (**c**) Feature quality evaluation by cluster compactness in latent space using Silhouette coefficient. (**d**) Spatial mapping and annotations of clusters with significant spatial co-localization patterns. Significantly co-localized clusters are identified using spatial co-localization score with permutation test. Clusters are assigned to one layer with respect to the anatomic annotations by original paper (**Methods**).

### Analysis of multi-modal clusters identified by MUSE

As a case study, we analyzed STARmap clusters identified from individual (based on PCA) or combined (based on MUSE) modalities in the joint latent space provided by MUSE (**Fig. 4a**). We classified clusters based on whether MUSE 1) refined, 2) reproduced or 3) discovered new clusters compared to those obtained from single modal analyses (**Supplementary Figs. 17** and **18**).

The **“refined”** MUSE clusters were poorly separated based on transcript features, yet were reasonably well separated based on morphology features (**Fig. 4b**, and **Supplementary Fig. 19**). In the combined analysis, MUSE employed the morphological diversity to further dissect cells into subgroups. Cell masks provided in this dataset are shown sampled randomly from each cluster, and morphological differences (e.g. small cell sizes of Cluster 11) can be seen (**Fig. 4b**, bottom). The **“reproduced”** MUSE clusters were distinct based on transcript features alone (**Fig. 4c**, top panel). Differential expression analysis allowed us to annotate these clusters as astrocytes, hippocampus neurons, oligodendrocytes or smooth muscle cell (SMC) types (**Fig. 4c**, bottom panel), which have distinct and identifiable transcriptional expression patterns. The **“discovered”** MUSE clusters were missed from the single modalities, which individually provided only weak differences (**Fig. 4d**, top panel). Here, the combination of weak heterogeneities from both modalities enabled MUSE to identify distinct L2/3, L5 and L6 structures (**Fig. 4d**, right bottom panel).

**Figure 4.**
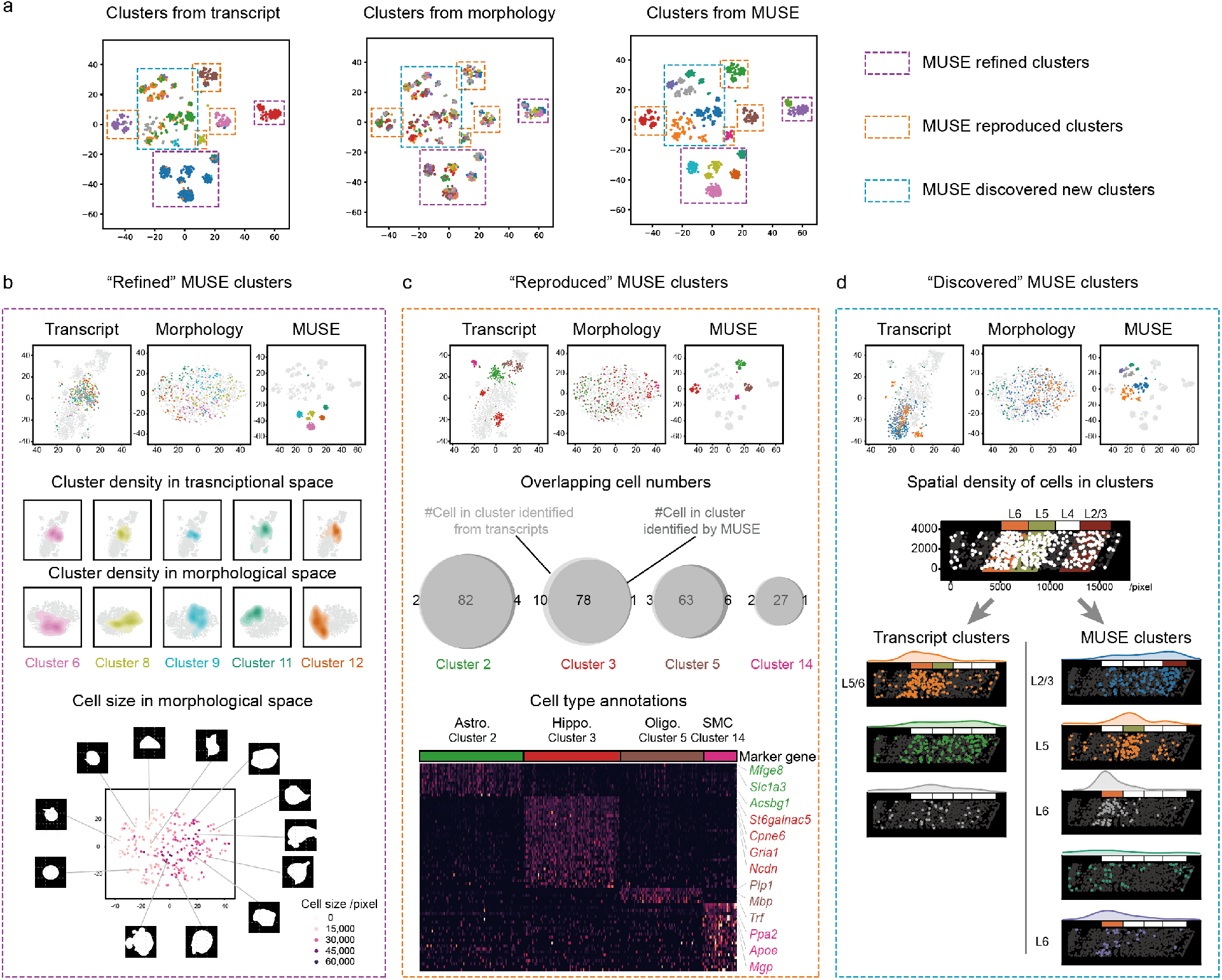
Analysis of MUSE clusters. (**a**) MUSE-identified clusters are categorized based on whether they refined, reproduced or discovered clusters compared to the single-modality PCA-identified clusters. tSNE visualization using MUSE latent space: cluster labels from transcript-only (PCA, left), morphology-only (PCA, middle) or combined (MUSE, right) analyses. (**b**) “Refined” MUSE clusters. Top: tSNE visualization of latent spaces. Middle: density plots of 5 MUSE clusters in transcriptional and morphological spaces using 2D kernel density estimation. Bottom: topography of cell size in tSNE representation of morphological space. Color: cell sizes. Points: cells in the 5 MUSE clusters. Cell masks: randomly selected cells. (**c**) “Reproduced” MUSE clusters. Top: tSNE visualization of latent spaces. Middle: Venn diagrams: number of overlapping cells between PCA transcript-only (grey outline) and MUSE (black outline) clusters. Bottom: “Reproduced” MUSE clusters identify astrocyte (Astro.), hippocampus neurons (Hippo.), oligodendrocyte (Oligo.) and smooth muscle cell (SMC) types (**Methods**). (**d**) “Discovered” MUSE clusters. Top: tSNE visualization of latent spaces. Bottom: “Discovered” MUSE clusters tend to be more layer specific than transcript-only clusters (density maps above tissue representations).

## Discussion

Characterizing cell heterogeneity is fundamental to understanding how tissues are organized and function. Two widely used and well-validated methods to study cell diversity are microscopy, to capture morphological differences, and scRNA sequencing, to capture transcriptional differences. Here, we developed a deep-learning framework, MUSE, to combine observations from both single-cell morphology and transcript modalities. MUSE makes use of a learning architecture that encourages synthesis of subpopulation structure observed in either modality. We demonstrated, for both synthetic and real biological data, that MUSE can reveal novel subpopulation structure and tissue organization missed by single-modalities or other methods.

It is evident that combined analysis across-omics modalities will increase our power to understand tissue heterogeneity^35, 36^. The machine learning approach of MUSE – with its parallelized autoencoder architecture and self-supervised learning approach – is designed to be extensible across modalities. MUSE is posed to leverage and coalesce advances as new measurement modalities, deeper profiling approaches, and modality-specific analysis are developed.

## Methods

### Multi-modal structured embedding

MUSE learns joint latent features by incorporating heterogeneity of morphological and transcriptional modalities. For a single-cell spatial transcriptomics dataset with *n* cells, transcriptional and morphological profiles are represented as *X* ∈ ℝ^*n×p*^ and *Y* ∈ ℝ^*n×q*^, where the *i*^th^ row of each matrix is the transcriptional (*x_i_*) or morphological (*y_i_*) feature from the same cell *i*.

#### Zero-inflated multi-modal autoencoder

The whole autoencoder structure is illustrated in **Supplementary Fig. 20**. Features from two modalities (*x_i_* and *y_i_*) are input into a multi-modal autoencoder, and a latent representation for each modality is learned by the encoder layer:

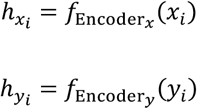

where 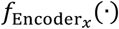, 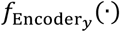 are multi-layer neural networks for two modalities and 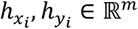, are latent representations with the same low dimension extracted from high-dimensional original inputs. The activation function in the last layer of the two encoders is chosen as tanh(·) to ensure the same scale for the two representations. Then the initial joint representation *z_i_* is learned by combing 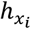 and 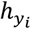:

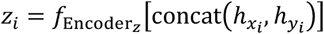

where concat(·) function concatenates two latent representations into one vector and the neural network encoder 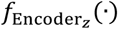 further encodes the vector into a joint representation *z_i_* ∈ ℝ^*k*^. The joint representation *z_i_* will be optimized by structured self-supervised loss.

Next, we use sparse weight matrices *w_x_* and *w_y_* to selectively activate entries in *z_i_* for the reconstruction of the original features *x_i_* and *y_i_*:

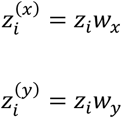

where *w_x_*, *w_y_* ∈ ℝ^*kxk*^ are sparse matrices that only employ a subset of the entries in *z_i_* to generate modality-specific features 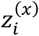 and 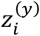. Finally, features for each modality are reconstructed by decoders:

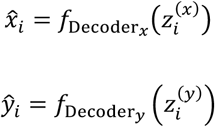

where 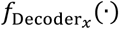 and 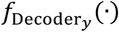 are multi-layer neural networks that expand latent representations into reconstructed features 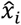 and 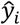.

#### Self-reconstruction loss

*For the transcriptional modality*, dropout is a major limitation due to the challenges of tracking fluorescent spots across multiple imaging rounds. Therefore, transcript profiles from *in situ* sequencing usually include a large proportion of zeros. Here we use a zero-inflated reconstruction error for the transcript modality to remove the effects of zeros entries:

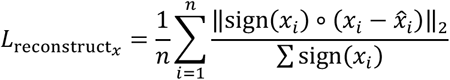

where sign(·) is a sign function that returns either 0 or 1 based on whether *x_i_* is zero or not (respectively); ∘ is the Hadamard product that conducts element-wise product of two vectors. The loss function calculates reconstruction errors for non-zero expression then averages them over non-zero entry numbers (Σ sign(*x_i_*)). *For the morphological modality,* we used the standard reconstruction loss:

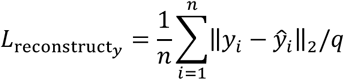

*The overall reconstruction loss* is the combination of the two modality losses with the sparsity constraint:

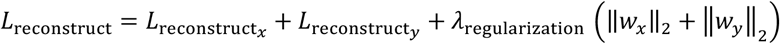

where *λ*_regularization_ is the regularization hyperparameter and is determined through analysis on simulation data (**Supplementary Fig. 21a**).

#### Structured self-supervised loss

Feature properties, noise levels and distributions, dropout levels are intrinsically different between the image and transcript modalities. To extract useful information from each modality and increase the quality of the joint latent feature *z_i_*, we further used structured self-supervised learning to encourage the structure of each modality to be maintained in the joint latent space.

To identify modality-specific population structure, clustering is performed for cell *i* using the latent feature from each modality 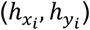 independently. Here, PhenoGraph^20^ was used to identify sample structures. The optimal cluster number is determined automatically on sample graph structures, an approach that is widely used in single-cell analysis. Here, cluster labels for cell *i* with respect to modality features *x_i_* and *y_i_* are denoted by 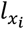 and 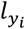, respectively. Cells with the same labels are similar to each other in (at least) one modality. Clusters from each modality are used as supervising labels to improve the learning of joint latent feature *z_i_* via the triplet loss:

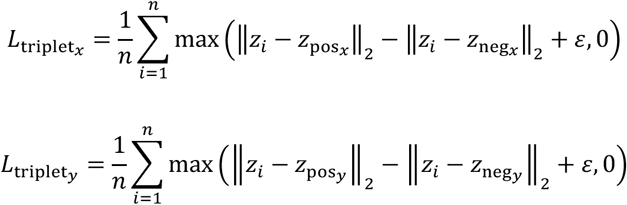

where in 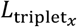, sample *z_i_* is the anchor, 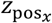 is a positive sample from the same cluster as the anchor based on clusters from modality 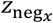 is a negative sample from a different cluster; *ε* is the margin; 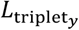 is defined in the same way using clusters from modality *y*. The triplet loss pushes the distance difference between anchor-positive and -negative samples to be greater than the margin so that the loss approaches its minimum (i.e. 0). As the choice of margin *ε* is hard to predetermine due to the uncertainty of feature distributions in two modalities, an adaptive method was used to automatically determine the margin value (refer to Optimization of MUSE).

#### Loss function

The overall loss function for training is the combination of the self-reconstruction and self-supervised losses:

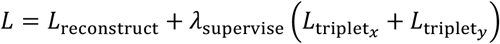

where *λ*_supervise_ is the hyperparameter to balance the contribution from triplet loss terms and was determined by simulation experiments (**Supplementary Fig. 21b**).

#### Optimization of MUSE

MUSE is trained on raw features and reference labels from two modality and optimizes joint latent features and cluster labels iteratively.

First, we obtain an estimate of the margin *ε* used in triplet loss. To accomplish this, we train the model without supervised terms by setting *λ*_supervise_ = 0, which is equivalent to a multi-modal autoencoder with zero-inflated loss in the transcript modality. We then estimate *ε* as the differences between medians in the top and bottom 20% values in the pairwise distance matrix from the initialized joint latent *z_i_*.

Then, we optimize the whole MUSE model using iterative training (over the complete loss function):

1. Fixing the network parameters, update the cluster labels 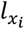 and 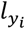 by using clustering on 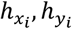, (see below).
2. Fixing cluster labels 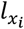 and 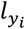, optimize the network parameters to obtain updated 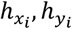, and *z_i_*.

#### Clustering

During training, clustering and labels for each independent modality were obtained using PhenoGraph^20^ with the Louvain method, which determines optimal cluster number automatically. After optimization, the same procedure was used to obtain clusters and labels for the joint latent space. For all PhenoGraph analysis, we used the same default 30 nearest neighbors to construct graph. We note that the architecture of MUSE is flexible, and other (e.g. modality-specialized) clustering approaches can be used instead of PhenoGraph to provide cluster labels 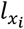 and 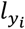.

### Spatial transcriptomics data preprocessing

We made use of two cortex datasets from seqFISH+^12^ and STARmap^15^, respectively. The seqFISH+ data includes 523 cells from 5 fields of views in a mouse cortex. For each cell, 10,000 RNAs were profiled using *in situ* sequencing. DAPI and Nissl stains were used in imaging, and cells were segmented manually based on their morphology. The STARmap dataset mapped 973 cells for mouse visual cortex, and each cell has 1,020 gene measurements. Cells were identified using watershed segmentation and segmentation masks were provided.

#### Transcriptional analysis

For both datasets, we performed preprocessing based on gene count data. We selected the top 500 most variable genes (genes with zero counts for all cells were excluded by default). Gene counts were normalized by library size and transformed using log(1+x) before input into the tested models. Transcript analyses were performed using *scanpy* python package (version 1.4.4)^37^.

#### Morphological analysis

For seqFISH+ data, we segmented single-cell images from tissue images for each imaging channel independently using the provided segmentation masks. Each cell-segmentation region was placed at the center of an empty image with 299×299 pixels. Then, DAPI and Nissl channels were independently input into the pretrained Inception v3 deep neural network. The output of the last network layer (with 1,024 dimensions) from each cell was concatenated into a long vector. PCA was applied to compress these feature vectors to 500-dimensional feature vectors. Vectors were scaled to have same mean value on all cells as in transcript features and then were used as input to the tested models. For STARmap data (which lacked cell markers), we directly placed the provided cell-segmentation masks over blank images then input them into the same pretrained neural network as for seqFISH+. Outputs from the last layers were also compressed using PCA and scaled to obtain single-cell morphological features. For both implementations, we used the Inception-v3 network^34^ with pretrained parameters provided by TensorFlow Hub (https://tfhub.dev/).

### Simulation experiment setup

We generated simulated ground-truth class labels *l* ∈ {1,…, *L*}^*n*^ for *n* cells and *L* possible cluster (i.e., cell subpopulation) assignments (see **Supplementary Table 1** for values of all parameters below). We simulated the situation for which only a proportion of true cluster identities could be observed from each modality, but all clusters could be discriminated using both modalities (**Supplementary Fig. 1**). To accomplish this, we divided the true clusters into two non-overlapping groups that were each assigned to one of the two modalities. Then, in each group, clusters were merged with probability *p* providing observed cluster labels *l_x_*, *l_y_* for the two modalities.

For example, 10 ground-truth clusters, labeled {1,…,10}, could be divided into groups *G*_1_ = {1,…,5}, *G*_1_ = {6,…,10} with modality 1 considering merges from *G*_1_ but not *G*_1_, and *vice versa* for modality 2; after merging, modality one might have seven clusters formed clusters {1 ⋅ 2 ⋅ 3, 4 ⋅ 5,6,7,8,9,10} while modality two might have six clusters formed from {1,2,3,4,5,6 ⋅ 7 ⋅ 8 ⋅ 9 ⋅ 10} (where " ⋅ " indicates merged clusters). While each modality can only distinguish a subset of the clusters, the combination has the potential to distinguish all of them.

For the transcriptional modality, we followed the same scRNA-seq simulation framework as used in SIMILR^26^ and scScope^27^. In short, we generated latent codes 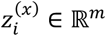 for cell *i* using a multivariable normal (MVN) distribution:

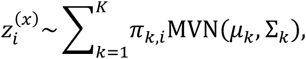

where *K* is the total cluster number; *π_k,i_* = 1 if cell *i* was assigned to cluster *k* in *l_x_* and otherwise 0; *μ_k_* ∈ ℝ^*m*^was sampled from a uniform distribution with Σ_*k*_ ∈ ℝ^*mxm*^ the identity matrix. Raw transcriptional features were generated through a linear transformation by 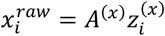, where entries in the random projection matrix *A*(^*x*^) ∈ ℝ^*pxm*^ were randomly sampled from the uniform distribution between [−0.5,0.5]. Gaussian noise was added to features 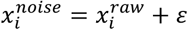, where *ε* was sampled from a Gaussian distribution *N*(0, *σ*^2^). Next, dropout in the count matrix with dropout rate proportional to expression level was simulated as:

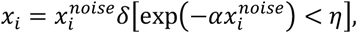

where δ[·] is an indicator function that outputs 1 if the argument is true and otherwise 0; *α* is the decay coefficient that controls dropout levels (set by default to 0.5); *η* is a random value sampled from the uniform distribution between [0, 1]. We input *x_i_* to all methods for analysis.

For the morphological modality, we generated latent codes 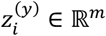 using the same mixture model procedure as above with modality labels *l_y_*. To add complexity to these “image-based” features, we passed these latent codes through a two-layer, non-linear network:

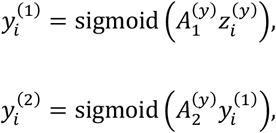

where 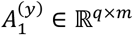 and 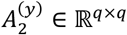 were matrices ran(domly sampled from the uniform distribution [−0.5, 0.5]; sigmoid(·) is the sigmoid function to non-linearly transform the data. Finally, as above we added random noise and dropouts to 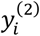 to obtain final morphological features *y_i_*. As a heuristic, the number of dropouts in this modality was set to 0.1 in order to obtain reasonably similar ARI scores for clustering based on each modality alone.

### Analysis of gene expression data

#### Differential analysis of expression data

With cluster labels, we identified differentially expressed genes using fold changes and p-values. For each cluster, we compared within *vs*. across cluster gene expression of cells. Log2 Fold changes were calculated based on mean gene expressions of these two groups to reveal the average expression differences. P-values were derived from one-sided ranksum test on expression profiles between the two groups to measure overall expression distribution differences.

#### Annotation of cortex layers

For the seqFISH+ data, marker genes that identify different cortex layers were obtained from the literature^32^ (in our case, four different genes were used to identify four different layers; **Supplementary Fig. 11**). Next, clusters with layer-like structures were identified (see below for score). Finally, for each layer-like cluster, the maximally overexpressed marker gene was used to assign each cluster to a layer.

For the STARmap data, anatomic layer labels were provided^15^. This allowed clusters to be annotated based on their spatial positions in the tissue section. First, clusters with significant spatial colocalization patterns

(based on spatial colocalization score) were identified for annotation. Next, a 1-dimensional kernel density estimation (KDE) with Gaussian kernels was performed along the provided x-coordinate of the image (corresponding to the cortex axis) for each cluster to model the spatial density of cells in the tissue. Finally, clusters were assigned to anatomic layers where peaks of cell spatial densities were located. In our implementation, kernel density estimation was performed using the KernelDensity function from sklearn python library with bandwidth determined by Scott’s rule.

## Evaluation of identified subpopulations

### Spatial co-localization score and evaluation

To quantify the spatial enrichment in the tissue for cell clusters, we designed a spatial co-localization score based on the statistic used in gene-set enrichment analysis (GSEA)^38^.

For all cells, we first calculated the cell-cell distance matrix *D* = {*d_ij_*} ∈ ℝ^*n×n*^, where *d_ij_* is the Euclidean distance between cells *i* and *j* on the image. The distance matrix was further converted into similarity *R* = {*r_ij_*} ∈ ℝ^*n×n*^ by taking *r_ij_* = 1/*d_ij_*, *i* ≠ *j*. As the similarity matrix is symmetric, only one similarity score is used for each cell pair (*i* < *j*). All off-diagonal, upper-triangle entries (*r_ij_*, *i* < *j*) in *R* were ordered into a list and re-indexed by rank *L* = {*r_k_*}, where *r_k_* is the similarity score in position *k* of *L*. If *n* is the total number of cells, then the size of *L* is given by *N* = (*n* − 1)(*n* − 2)/2.

We define two scores that allow us to assess whether a cluster label, *C*, is consistent with distance similarities. First, let *S_c_* ⊂ *L* be the set of similarity scores *r_k_* obtained from cells within *C* and define:

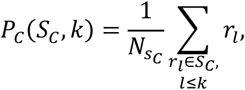

where 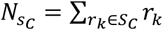. Second, for *r_k_* ∉ *S_C_* (i.e., at least one cell is not in *C*), define:

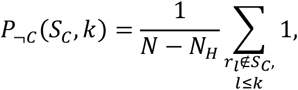

where *N_C_* = (*n_c_* − 1)(*n_c_* − 2)/2 is the size of *S_c_* (*n_c_* is the number of cells in *C*). The spatial co-localization score (SCS) for *C* is defined as the maximal signed deviation between distributions *P_C_*(*S_c_*, ·) and 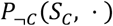.

To derive a significance p-value, we constructed null SCS distribution by permuting cluster labels and calculating corresponding scores 1000 times. The p-value is defined as the proportion of scores greater than SCS on non-permuted labels. Source code for the SCS calculation is provide on GitHub.

#### Cluster accuracy evaluation with adjusted Rand index

For simulation studies, where ground truth subpopulation labels were given, we evaluated clustering performance using the adjusted Rand index (ARI)^25^. An ARI near 1 indicates a strong match to ground truth clustering, while values near 0 suggest random assignment. In the implementation, we used the adjusted_rand_score function from sklearn.metrics python package.

#### Feature quality evaluation with Silhouette Coefficient

The quality of latent features were evaluated by the compactness of the clusters in the latent space using the Silhouette coefficient^39^. A score of 1 indicates highest density in latent space. In our implementation, we employed the silhouette_score function from sklearn.metrics python package.

### Comparing methods

All compared methods were run on the same input features (see above data processing section; single-modal methods took features from only one modality) to learn 100-dimensional latent representations. Subpopulations were identified based on latent representations using PhenGraph^20^. All methods were configured with default parameters (unless specifically noted) and were run on the same Linux desktop (Ubuntu 18.04.3 LTS operation system) with Xeon E5 CPU and Nvidia Titan X GPU (Driver Version: 418.87.00, CUDA Version: 10.1).

The software packages used for comparisons are as follows.

#### Transcriptional feature learning methods

Principle component analysis (PCA): sklearn 0.20.3 python package. Zero-Inflated Factor Analysis (ZIFA)^40^: ZIFA.fitModel() from ZIFA python package (v0.1). Single-cell Interpretation via Multi-kernel LeaRning (SIMLR)^26^: python implementation of SIMLR (v0.1.3). scScope^27^: python implementation scScope (v0.1.5).

#### Morphological feature learning methods

PCA: as above. Multi-dimensional scaling (MDS), Isometric mapping (Isomap), and t-distributed stochastic neighbor embedding (tSNE): sklearn.manifold python library (v0.20.3). We note that the tSNE method can only support maximal 3-dimension latent representations.

#### Multi-modal feature learning methods

Correlation analysis (CCA) learns linear transformations of multiview data and maximizes their correlations in latent spaces, and we chose the transformation of the transcript data for clustering: sklearn python package (v0.20.3). Multi-omics factor analysis v2 (MOFA+)^24^ was designed to combine multi-omics data using the multiview matrix factorization: mofapy2 package (v0.3) with factors = 100, iteration = 500, group number = 1 and view number = 2 to learn 100-dimension joint features. Autoencoder (AE) learns joint representations based on reconstruction loss from two modal features: in the implementation, we used the same neural network structure as in MUSE with standard reconstruction loss with all learning parameters (learning step, iteration numbers, etc.) the same as used in MUSE. Multi-modal structured embedding (MUSE): software is implemented in python 3.7.3 with NumPy, SciPy, PhenoGraph and TensorFlow packages (details were provided at https://github.com/AltschulerWu-Lab/MUSE); hyperparameter values were chosen through simulation study (**Supplementary Fig. 21**) and are provided in **Supplementary Table 2**.

### Software used in the study

Software packages use in the study can be accessed *via* following links PhenoGraph: https://github.com/jacoblevine/PhenoGraph

ZIFA: https://github.com/epierson9/ZIFA

SIMLR: https://github.com/bowang87/SIMLR_PY

scScope: https://github.com/AltschulerWu-Lab/scScope

MOFA+: https://github.com/bioFAM/MOFA2

seaborn: https://seaborn.pydata.org/

sklearn: https://scikit-learn.org/

## Data and code availability

### Single-cell spatial transcriptomics datasets

seqFISH+: transcript data were downloaded from the GitHub page of seqFISH+ project (date: August 1, 2019; link: https://github.com/CaiGroup/seqFISH-PLUS). Nissl and DAPI stained images were provided by authors of seqFISH+ paper.

STARmap: raw data were downloaded from the project page (https://www.starmapresources.com/data at July 2, 2019). Transcript profiles and cell segmentation masks were extracted from data using the python pipeline provided by authors at https://github.com/weallen/STARmap.

### Simulated tool for multi-modality data generation

Simulation code is available from GitHub https://github.com/AltschulerWu-Lab/MUSE.

### MUSE

MUSE is provided as a python package under MIT license and can be installed through “pip install muse_sc”. Source and demonstration code are available on GitHub https://github.com/AltschulerWu-Lab/MUSE.

## Acknowledgements

We thank Chee-Huat Linus Eng at Caltech for providing seqFISH+ image data, Xiao Wang at the Broad Institute and MIT for providing information on STARmap data analysis, and Olivier Moindrot at Stanford for the open-source implementation of the triplet loss. We thank Heinz Hammerlindl, Lee Rao, Susan Shen, Xiaoxiao Sun and other members of the Altschuler and Wu labs for constructive feedback. S.J.A. and L.F.W gratefully acknowledge support from the UCSF Program for Breakthrough Biomedical Research, ProjectALS and the CZI NDNC Challenge Network.

