## Supplementary Materials for "Characterizing tissue composition through combined analysis of single-cell morphologies and transcriptional states"

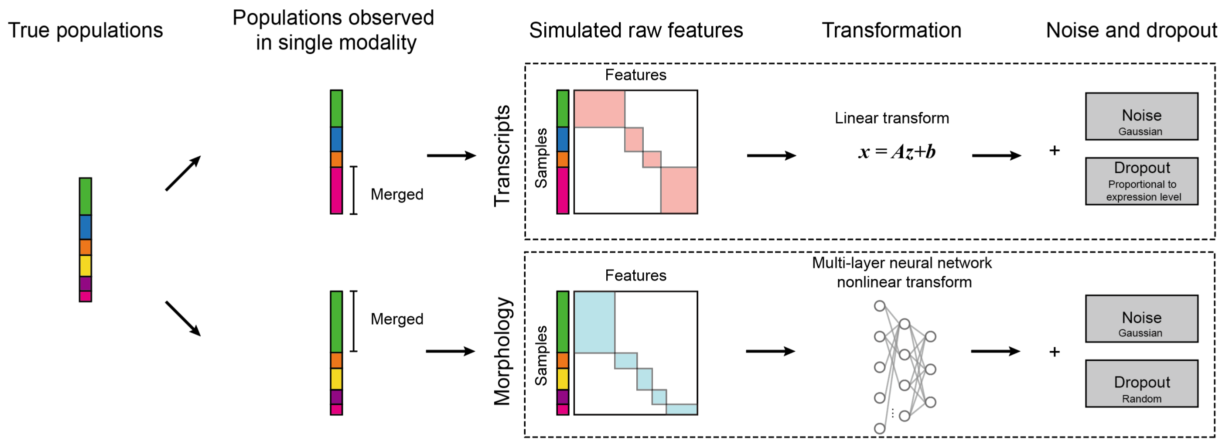


**Supplementary Figure 1 |** Simulation design to generate single-cell profiles with two modalities.

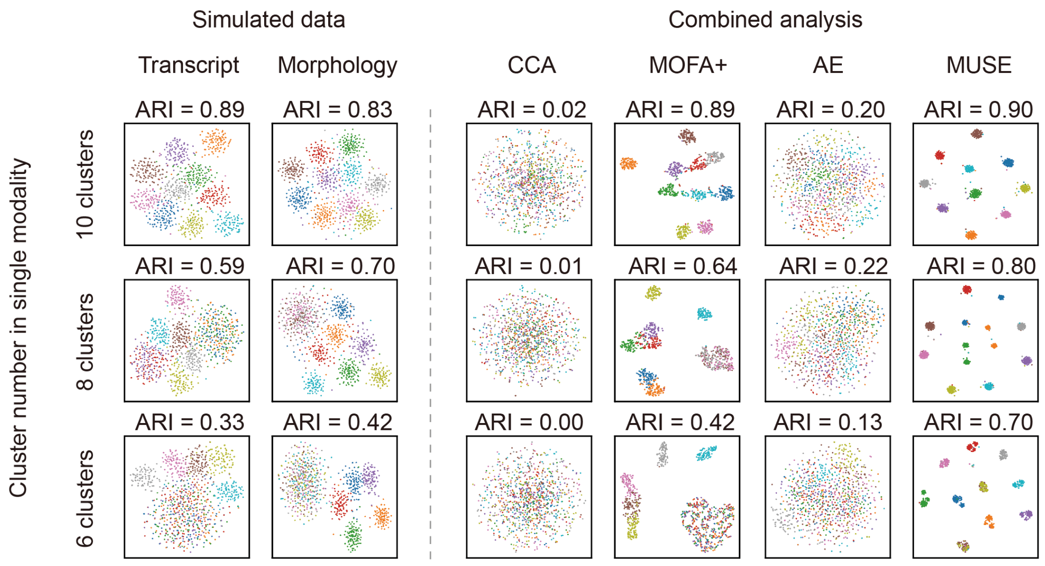


**Supplementary Figure 2 |** tSNE visualizations of latent representations from single- and combined-modality methods for randomly selected simulation experiments in **Fig. 1c**. Colors: ground truth subpopulation labels in simulation; n=1,000 cells; ground truth cluster: 10.

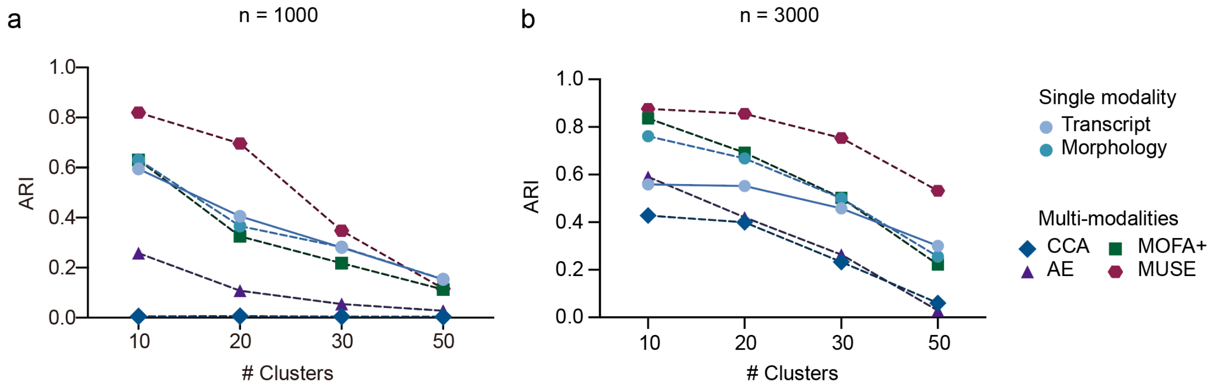


**Supplementary Figure 3 |** Evaluations of combined methods in simulated data with different ground truth cluster numbers. n = 1,000 (a) and 3,000 (b) samples were considered in simulations. Parameters used in simulation were listed in **Supplementary Table 1**.

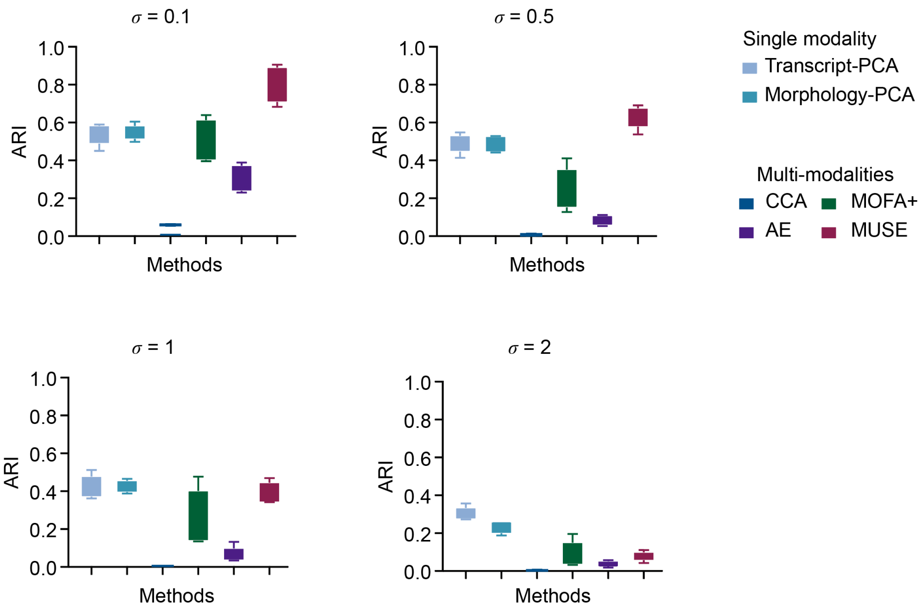


**Supplementary Figure 4** | Evaluations of combined methods in simulated data with Gaussian noise of various variance ($\sigma$). n = 1,000 cells; ground truth cluster: 10; other parameters used in simulation: see **Supplementary Table 1**.

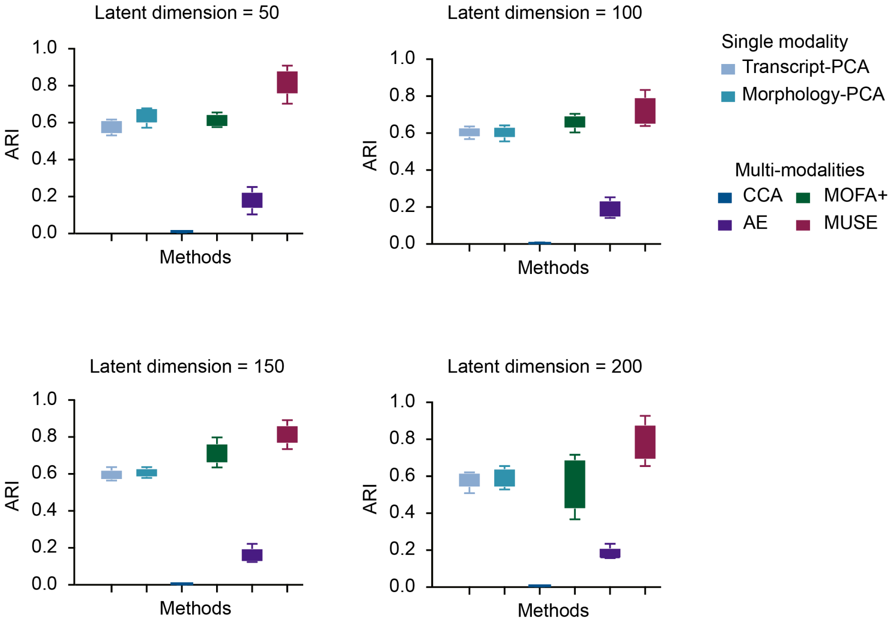


**Supplementary Figure 5 |** Evaluations of clustering accuracies under different dimensions of joint latent representations. n = 1,000 cells; ground truth cluster: 10.

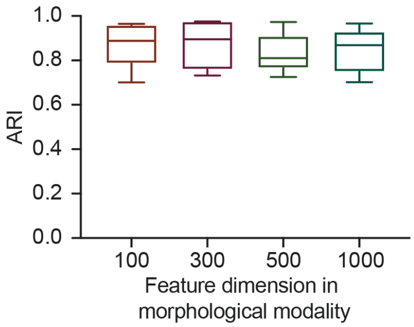


**Supplementary Figure 6 |** Clustering accuracies of MUSE when fixing the input dimension of transcript to 500 while changing dimension of morphological features between 100 to 1,000. n = 1,000 cells; ground truth cluster: 10; other parameters used in simulation: see **Supplementary Table 1**.

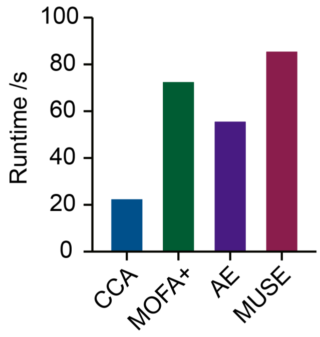


**Supplementary Figure 7 |** Run time of comparing methods on simulated data with 1,000 cells. We note for MOFA+, it has the GPU mode but we failed to configure it in our desktop.

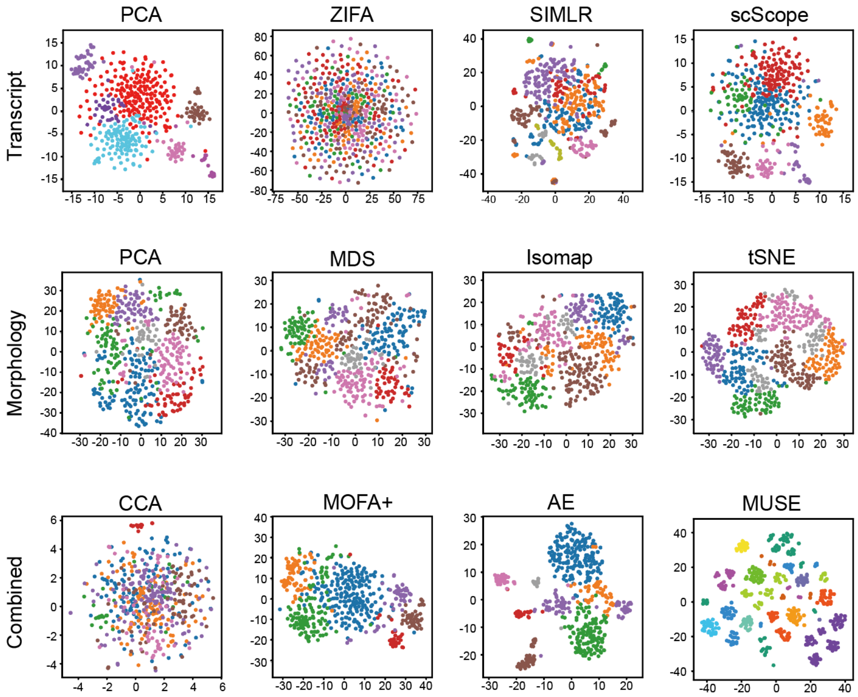


**Supplementary Figure 8 |** tSNE visualization of latent spaces from transcriptional (top), morphological (middle) and combined (bottom) methods on seqFISH+ data. Color: cell clusters identified from latent representations using PhenoGraph.

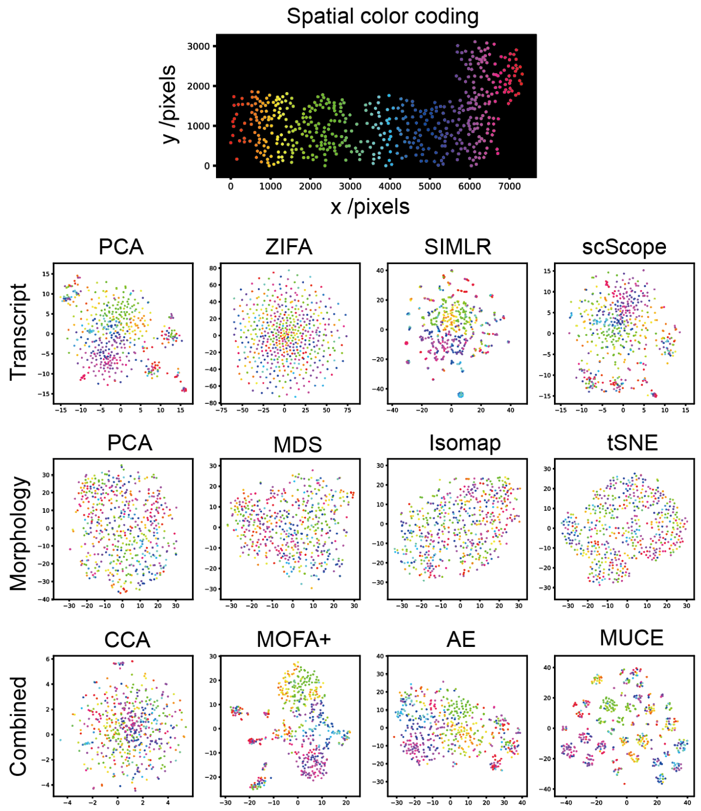


**Supplementary Figure 9 |** tSNE visualization of latent representations by different methods with pseudo-colors labeling cortex depth along x-coordinate for seqFISH+ data.

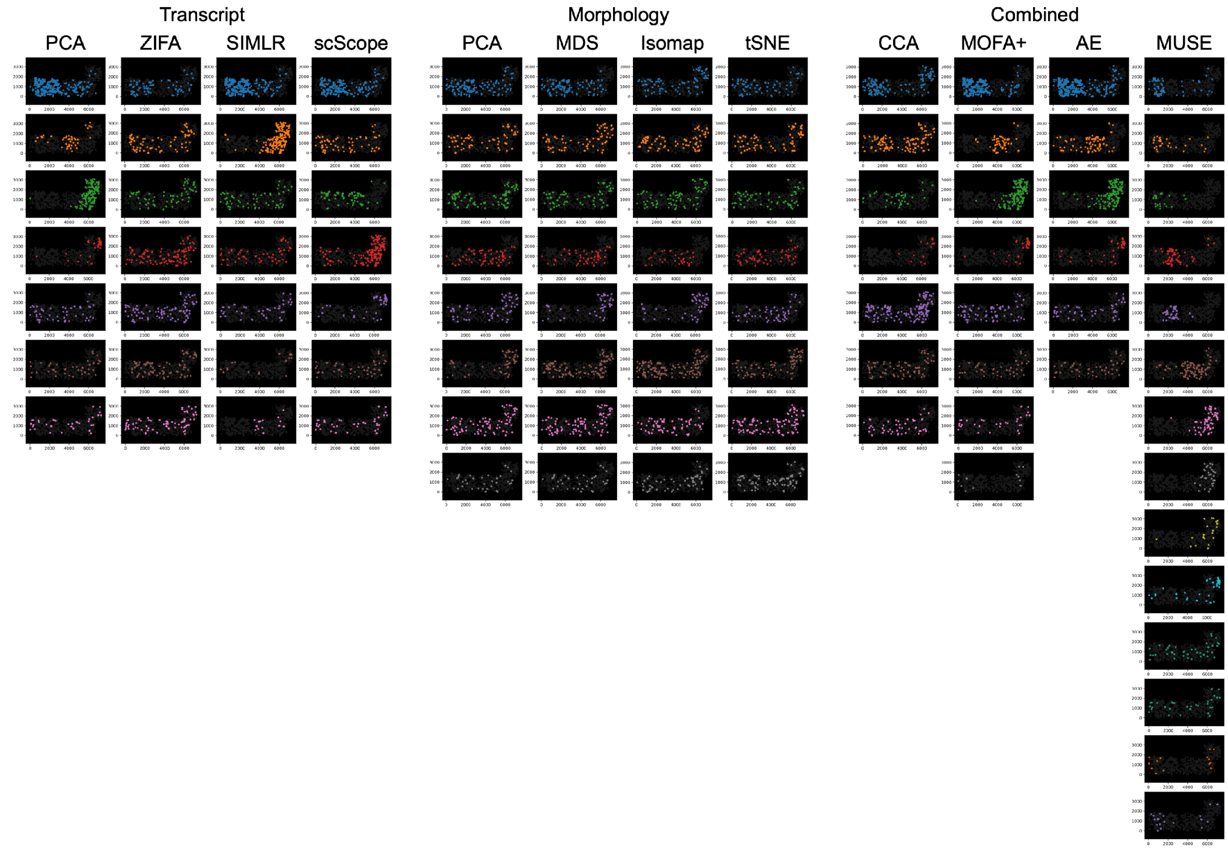


**Supplementary Figure 10** | Spatial mappings of cell clusters in the tissue section for seqFISH+ data.

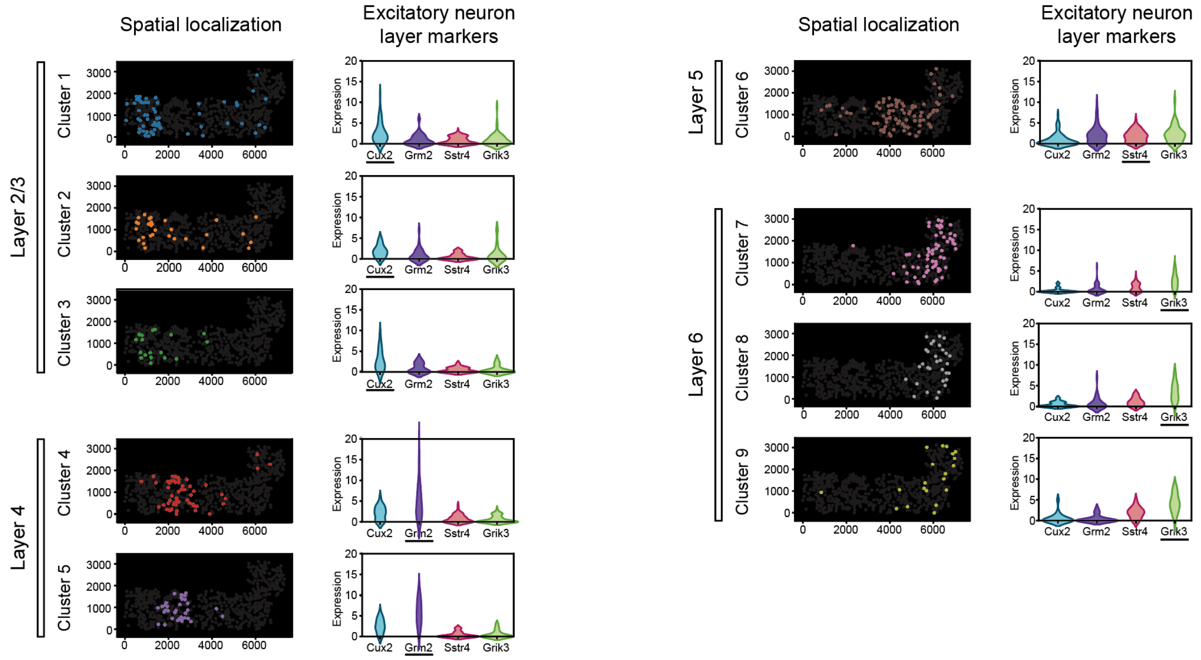


**Supplementary Figure 11** | Layer annotations of MUSE clusters based on layer gene markers for seqFISH+ data. Spatial localization of cell clusters (first column) and marker expression abundances (second column) were shown. For each cluster, gene names with maximal overexpression levels were underlined.

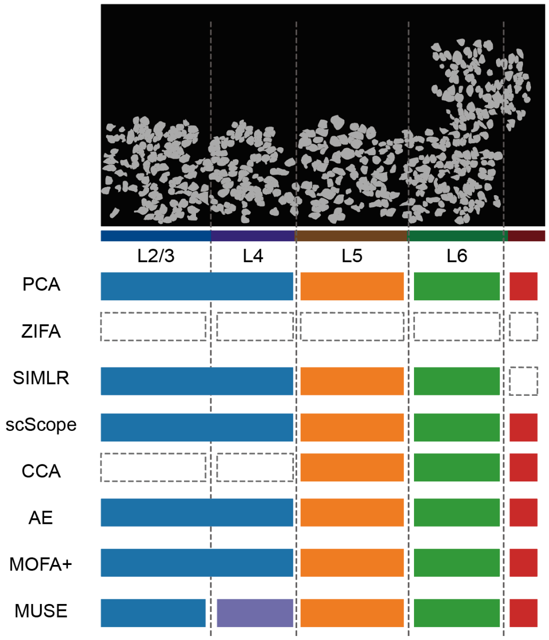


**Supplementary Figure 12 |** Comparison of discovered cortical layers by transcriptional or combined methods on seqFISH+ cortex data. 5 layers were shown. Squares with the same color and across multiple layers indicate the method discovered merged layers. Squares with no color indicate the method failed to discover the corresponding layer.

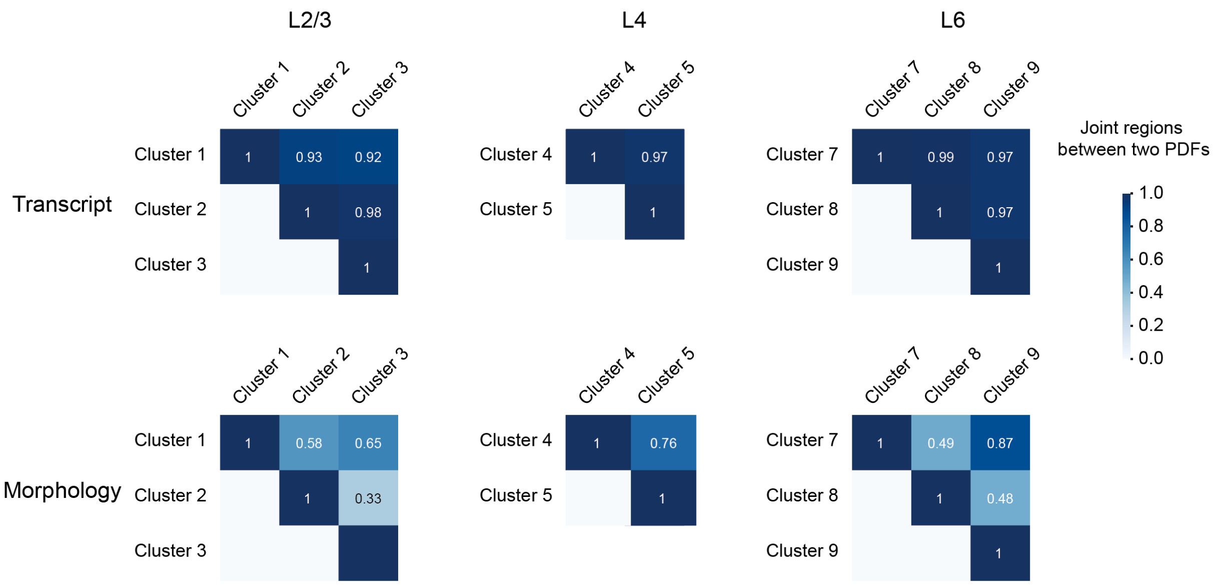


**Supplementary Figure 13 |** Overlapping areas of PC-1 distribution densities for transcriptional (top panel) or morphological (bottom panel) features, for MUSE clusters that are identified in the same cortical layers (corresponding to **Fig. 1e**).

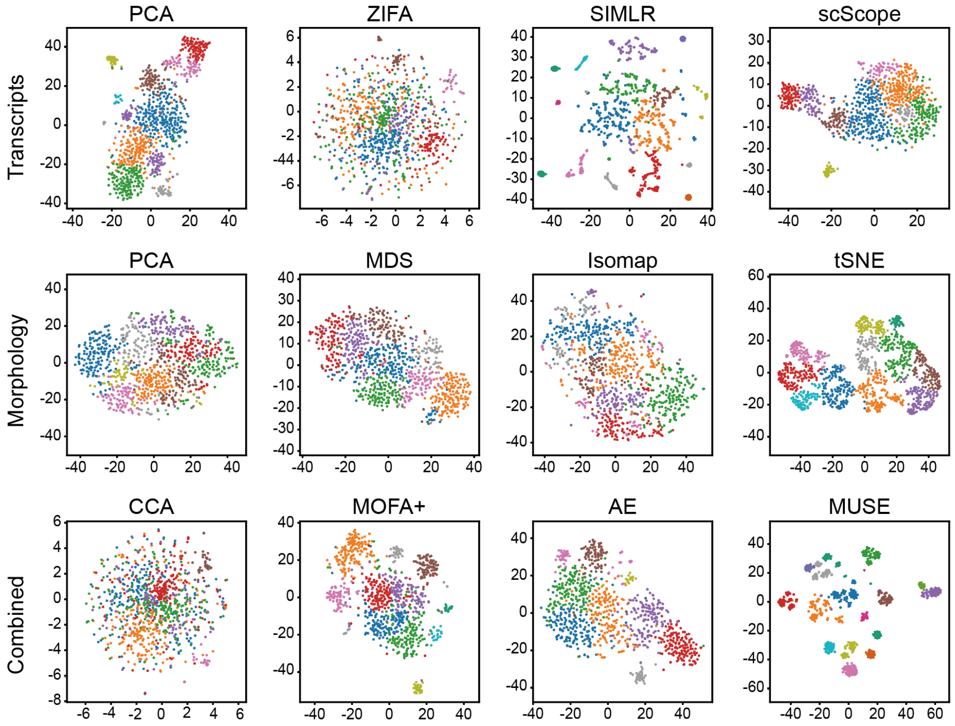


**Supplementary Figure 14 |** tSNE visualization of latent spaces from transcriptional (top), morphological (middle) and combined (bottom) methods on STARmap cortex data. Color: cell clusters identified from latent representations using PhenoGraph.

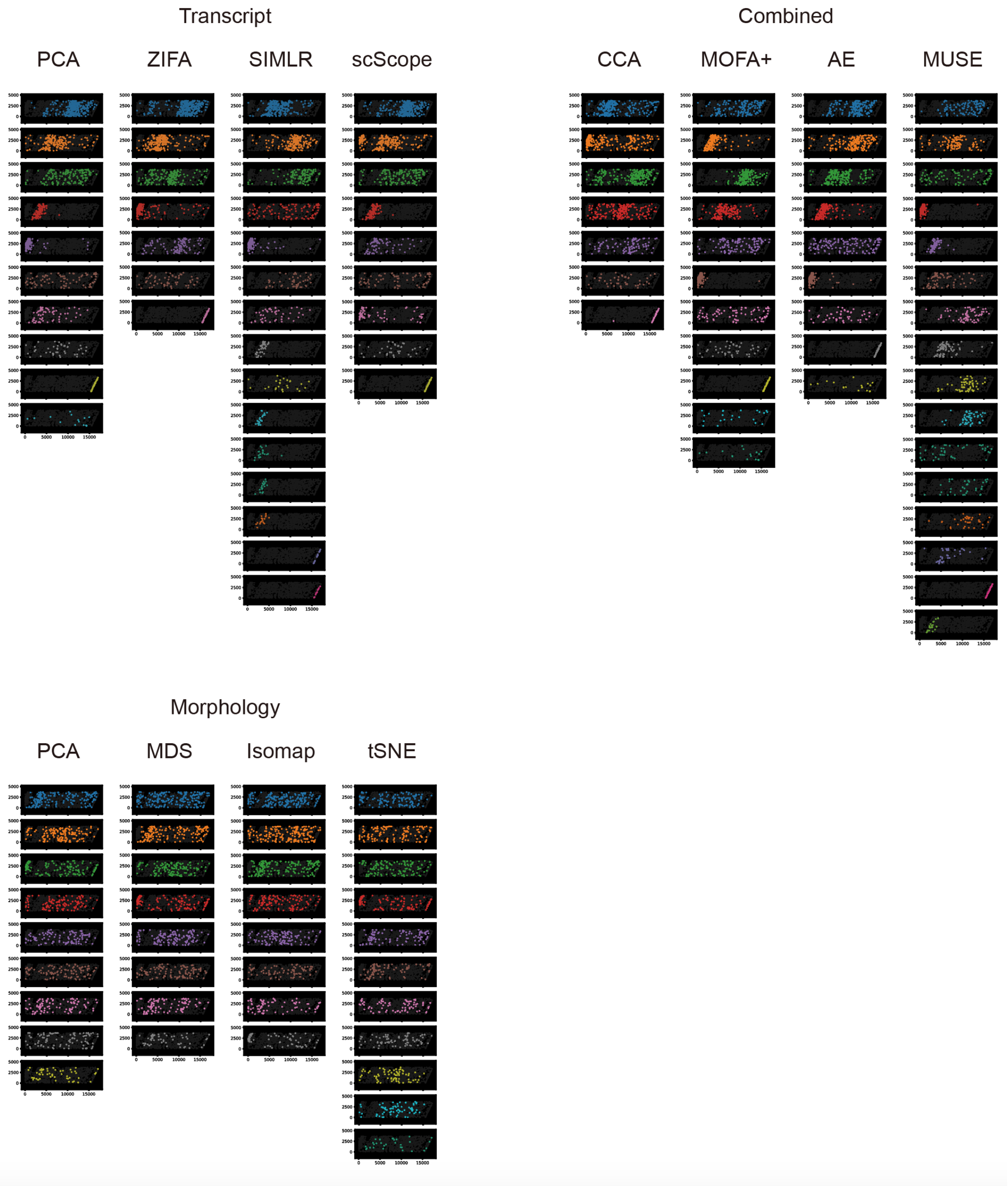


**Supplementary Figure 15 |** Spatial mappings of cell clusters in the tissue section for STARmap data.

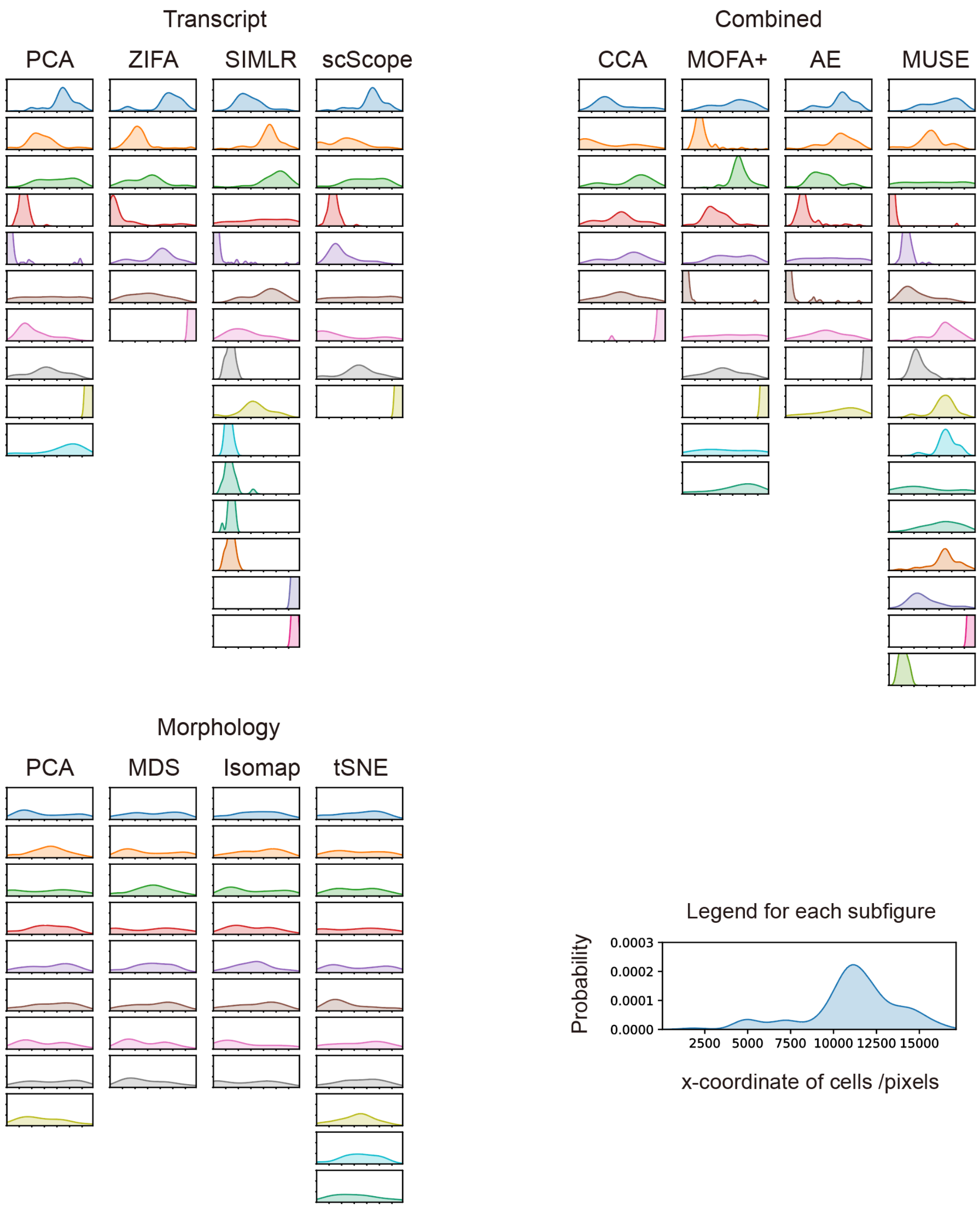


**Supplementary Figure 16 |** Spatial density plots in the tissue section for all clusters identified in STARmap data.

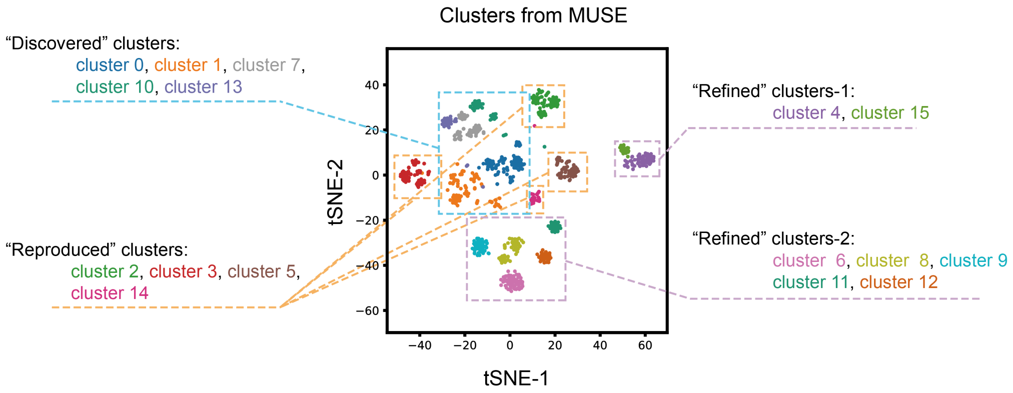


**Supplementary Figure 17 |** tSNE visualization of MUSE clusters in latent space for STARmap data. All clusters were classified into "Refined", "Reproduced" or "Discovered" types based on comparison with clusters identified from transcript-alone or morphological-alone analysis (corresponding to **Fig. 4a**).

a. “Refined” MUSE clusters b. “Repoduced” MUSE clusters c. “Discovered” MUSE clusters

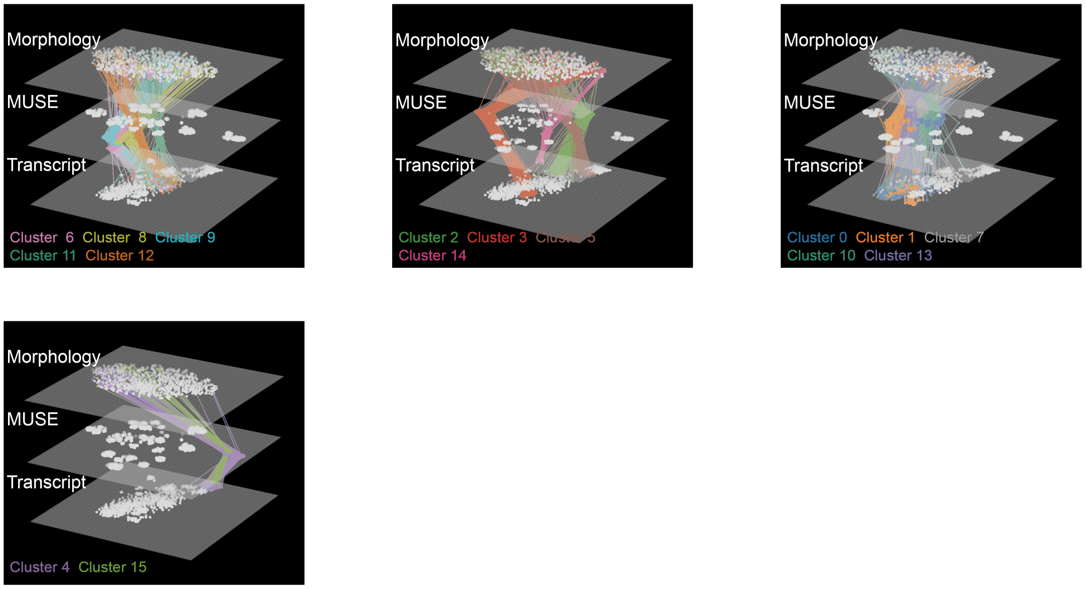


**Supplementary Figure 18 |** tSNE visualization of three types of MUSE clusters in the latent space of morphological features (top layer of each 3D plot), MUSE latent features (middle layer) or transcriptional features (bottom layer) for STARmap data. Same cells in three spaces were connected in lines.

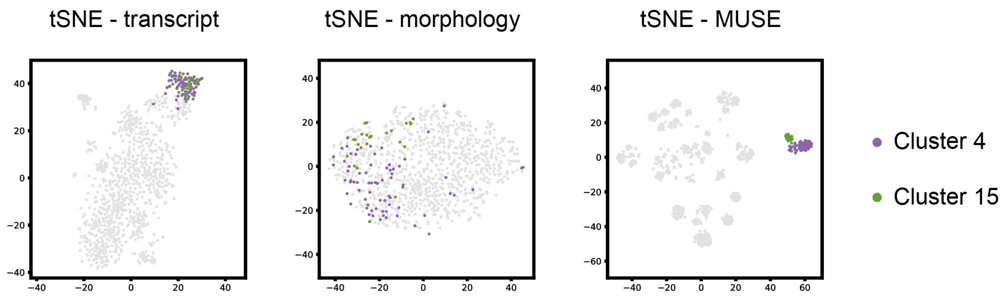


**Supplementary Figure 19 |** tSNE visualization “refined” clusters-2 in three latent spaces (related to **Fig. 4b**).

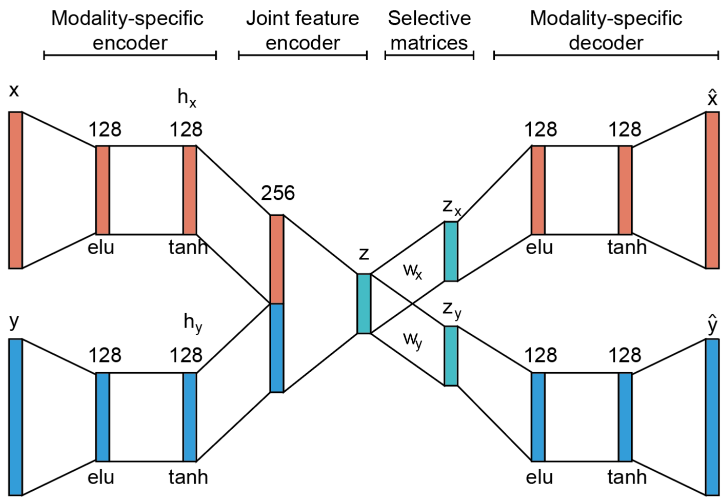


**Supplementary Figure 20 |** Model structure of zero-inflated multi-modal autoencoder used in MUSE.

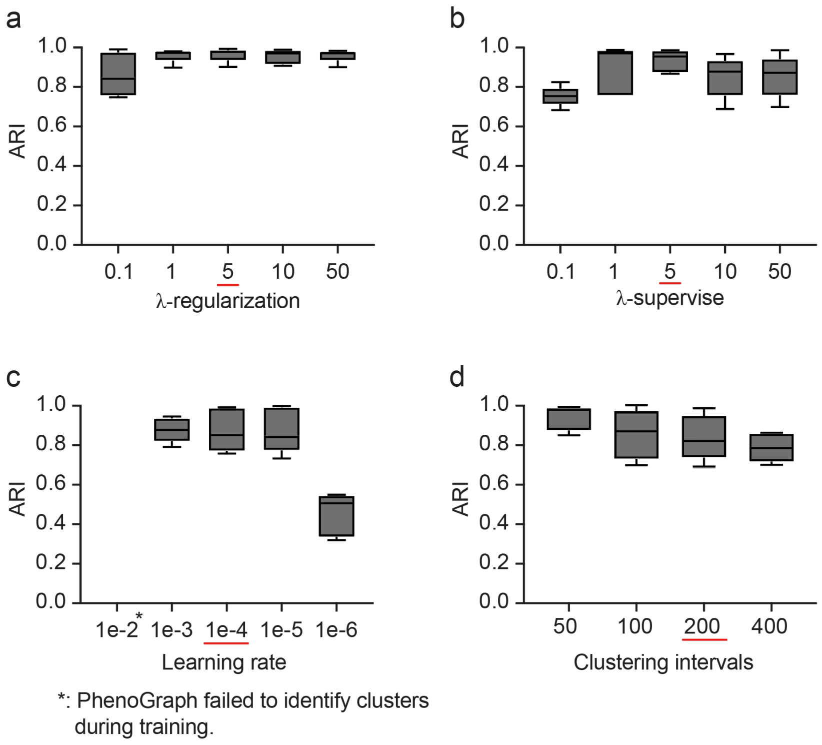


**Supplementary Figure 21 |** Performance evaluation of MUSE with different hyperparameter settings: (a) weight of regularization term; (b) weight of supervision term; (c) learning rate; and (d) iteration intervals between cluster updating in training. Red underlines: parameters selected as default in MUSE package.

**Supplementary Table 1** | Parameter settings used in simulation experiments.

| Experiment | Purpose | Ground truth cluster | Cluster merge probability | sample size | latent code dimension | Feature dimension | | Noise $\sigma$ | | Dropout coefficient | |
|---|---|---|---|---|---|---|---|---|---|---|---|
| | | | | | | Trans. | Morph. | Trans. | Morph. | Trans. | Morph. |
| **Fig. 1c Supplementary Fig. 2** | Performance when the ability to discriminate subpopulations in each modality decreases | 10 | Each single modality had 10, 8, or 6 clusters | 1,000 | 30 | 500 | 500 | 0.1 | 0.1 | 0.5 | 0.1 |
| **Figs. 1d, e** | Performance as data quality in one modality degrades by dropouts | 10 | 0.7 | 1,000 | 30 | 500 | 500 | 0.1 | 0.1 | 0.05, 0.1, 0.15, 0.2, 0.25, 0.3, 0.4, 0.5 | 0.1 |
| **Supplementary Fig. 3** | Performance as the number of ground-truth subpopulations increases | 10, 20, 30, 50 | 0.7 | a) 1,000 b) 3,000 | 30 | 500 | 500 | 0.1 | 0.1 | 0.5 | 0.1 |
| **Supplementary Fig. 4** | Performance with Gaussian random noise with increasing variance | 10 | 0.7 | 1,000 | 30 | 500 | 500 | 0.1, 0.5, 1, 2 | 0.1, 0.5, 1, 2 | 0.5 | 0.1 |
| **Supplementary Figure 6** | Effects from input dimensions | 10 | 0.7 | 1,000 | 30 | 500 | 100, 300, 500, 1,000 | 0.1 | 0.1 | 0.5 | 0.1 |

**Supplementary Table 2** | Default parameters used in MUSE.

| Hyperparameter | Value |
|---|---|
| Latent dimension | 100 |
| Learning rate | 1.00E-04 |
| Training epoch | 500 |
| Initialization epoch | 200 |
| Cluster update interval | 200 |
| λ_supervise | 5 |
| λ_regularization | 5 |
